# Temporal foundation model unveils ancient Hepatitis B virus evolution

**DOI:** 10.1101/2025.07.09.663843

**Authors:** Ji Qi, Bing Sun, Yonghao Liu, Xiaoming Xiao, Mi Zhou, Alexander Herbig, Chen Yang, Xiaosong Han, Yinqiu Cui, Renchu Guan

## Abstract

Since ancient hepatitis B virus (HBV) sequencing data are scarce and incomplete, the evolutionary dynamics of HBV have long remained enigmatic. This data gap hinders a comprehensive understanding of HBV genetic variations, evolutionary trajectories, and host interactions in different historical periods, preventing insight into its long-standing co-evolutionary relationship with humans. In this study, we developed a sequence-based generative foundational model, ViraChron (Viral Chronology Foundation Model), that integrates multi-scale temporal information to compensate for the lack of ancient HBV data. We confirmed the robustness of our model through sequence identity, phylogenetic, and recombination analyses. ViraChron enabled the identification of fixed mutations specific to HBV genotype A, demonstrating that HBV acquires survival advantages through mutations that are ultimately fixed by natural selection. Our model outperforms the original baseline in terms of sequence completion, achieving significantly higher accuracy. The model significantly reduced data acquisition time from 80 hours to 5.4 minutes per HBV genome, providing reliable, high-quality data for 7 new HBV genomes from specific archeological sites. ViraChron overcomes previous limitations, obtaining high-quality genomic data from samples over ten thousand years old, thereby facilitating a systematic understanding of viral evolution and its biological characteristics.

## Introduction

Evolution is fundamentally based on genetic changes that emerge and undergo selection over time[1]. Therefore, obtaining nucleotide sequence data from different time points represents the most direct and effective approach to studying evolutionary processes. Specifically, ancient sequences represent extinct genetic diversity, and these time-encoded sequences can fill in missing intermediate evolutionary lineages, making them crucial for reconstructing evolutionary pathways. However, analyzing data from ancient DNA poses inherent challenges, such as sample fragmentation and scarcity, which introduce uncertainties and reduce precision in key areas like molecular clock calibration and genome recombination analysis.

While current genomic language models perform exceptionally well with modern genomic data[2–7], they encounter significant difficulties when applied to ancient DNA reconstruction. Ancient genomic patterns are closely tied to temporal context, making it challenging for context-dependent inference models to accurately capture historical point mutations. Furthermore, missing fragments in ancient sequences may cause models to overlook critical genetic information or even prevent them from learning meaningful genomic patterns.

The hepatitis B virus (HBV), with its high mutation rate and long-term history of co-evolution with human hosts, serves as an ideal model for studying viral evolution. Recent rapid accumulation of ancient HBV data has provided unique opportunities to reveal its spatiotemporal spread and adaptive evolution[8–14]. Yet the fragmentary nature of ancient HBV data prevents systematic analysis of its evolutionary dynamics using conventional methods[15].

To address these challenges, we developed ViraChron (Viral Chronology Foundation Model), a novel framework that integrates multi-scale temporal encoding into the Evo model architecture. By transforming discrete temporal information into high-dimensional time-series tensors, our model successfully captures the dynamic evolutionary trajectories of viruses (Figure 1). Experimental results demonstrate that ViraChron not only generates complete HBV genomes corresponding to specific historical periods, but also accurately imputes missing sequences in genotypes A, B, C and D, with biological plausibility confirmed by Bayesian skyline analysis. This breakthrough establishes a new paradigm for overcoming limitations in analyzing ancient DNA data and significantly enhances the spatiotemporal resolution of evolutionary studies.

**Fig. 1.**
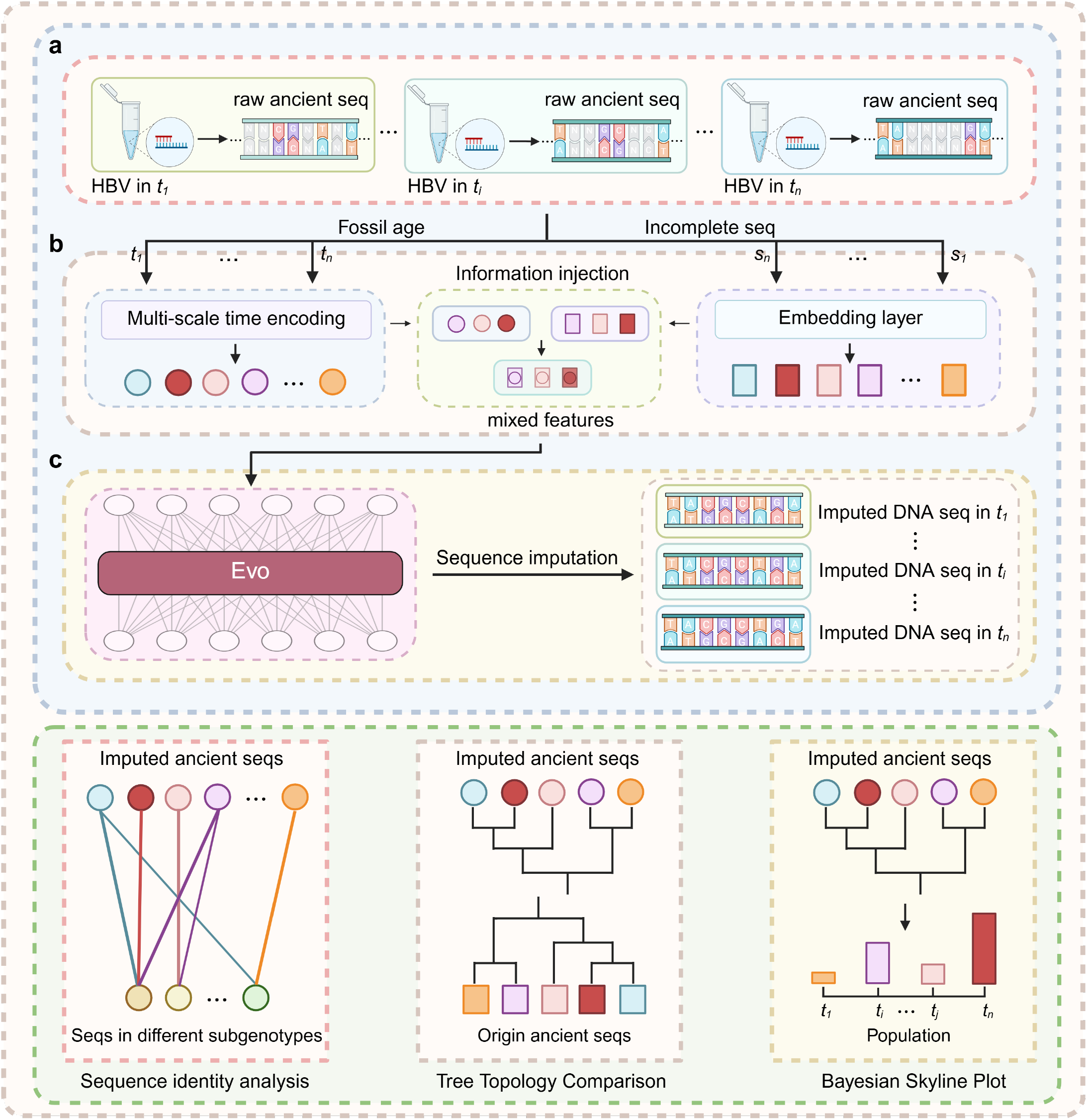
ViraChron Framework. (a) Ancient DNA genomes were captured to obtain the corresponding sequences and sampling times. (b) Sequences were embedded through the original embedding layer, while time was transformed into a high-dimensional representation using a time encoding module. The temporal information was then injected into the sequence embeddings. (c) We captured the evolutionary changes in HBV using Evo, ultimately reconstructing complete genomes. We recovered the raw DNA sequences and performed sequence identity analysis, phylogenetic analysis, and Bayesian skyline analysis (where the height of the bars represents population size). This figure was created in https://BioRender.com

## Results

### Ancient HBV DNA screening

We analyzed 68 sequencing datasets from different time periods to extract HBV DNA reads. Through this screening, we identified HBV-mapped reads in seven individuals from five archaeological sites dating roughly between 3000 and 1000 years ago (Figure 2 and Table S1). Five of these seven samples were from ancient Chinese sites, and two were from Russia. All individuals exhibited the characteristic damage patterns expected of ancient DNA (Figure S1). To improve the quality of the dataset, we performed in-solution capture enrichment targeting HBV DNA for all samples yielding HBV reads. Following enrichment, genomic sequences were reconstructed by mapping the reads to an HBV reference genome, resulting in coverage ranging from 78% to 100%, with an average depth of coverage between 11.6× and 495.2× (Table 1).

**Table 1.**
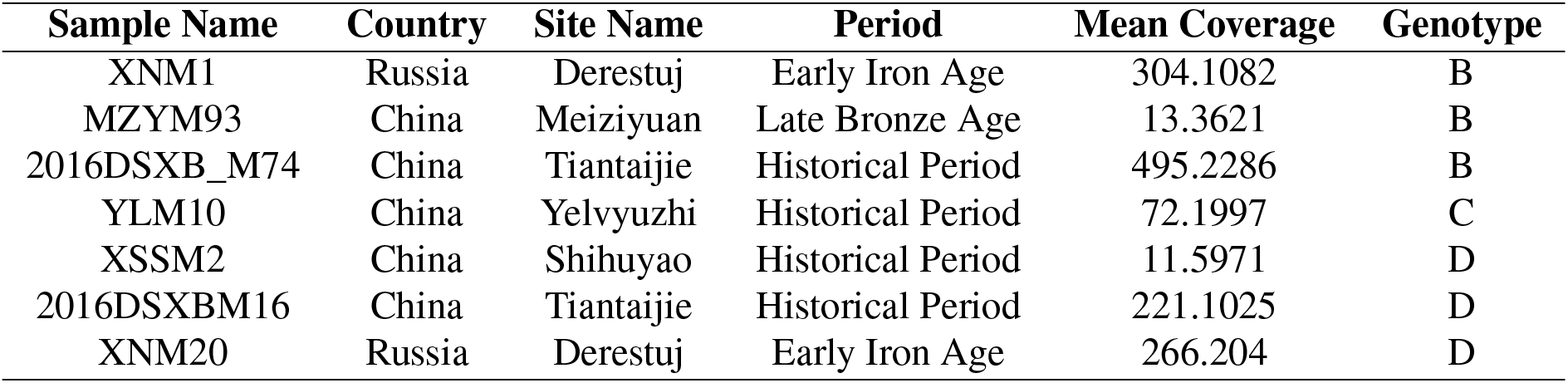
Summary of information for the individuals from which ancient HBV genomes were recovered in this study.

**Fig. 2.**
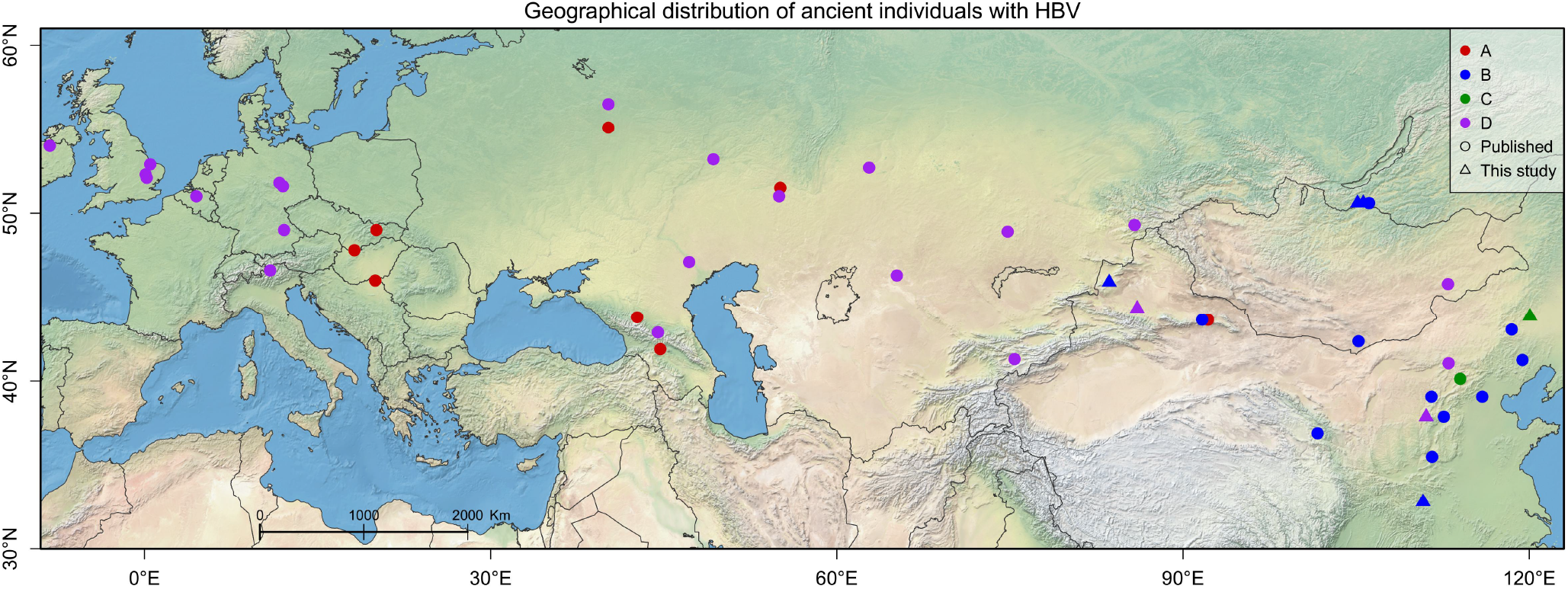
Geographical distribution of ancient individuals with HBV. Different colors represent different genotypes. Published datasets were utilized for model training, while newly acquired data from this study served as the test set.

### ViraChron learns the patterns of HBV genomes

To learn and represent patterns in HBV genomes such as genome organization, mutation hotspots, recombination and evolutionary dynamics, we collected 7804 complete and highconfidence HBV genome sequences from NCBI. After data alignment and cleaning, we built a dataset consisting of 7,804 ancient and modern HBV genome sequences for model training (Methods). We adopted a parallel fine-tuning strategy using both ancient and modern data, since fine-tuning on exclusively modern or ancient data tends to result in large divergence from the established evolutionary trajectories (Figure 3a). To determine the genotypes of the newly recovered ancient HBV genomes, we performed competitive mapping against representative genomes from each known lineage, following previously established protocols4. Based on the mapping results, we classified the seven ancient HBV genomes into three distinct genotypes (Table S1).

**Fig. 3.**
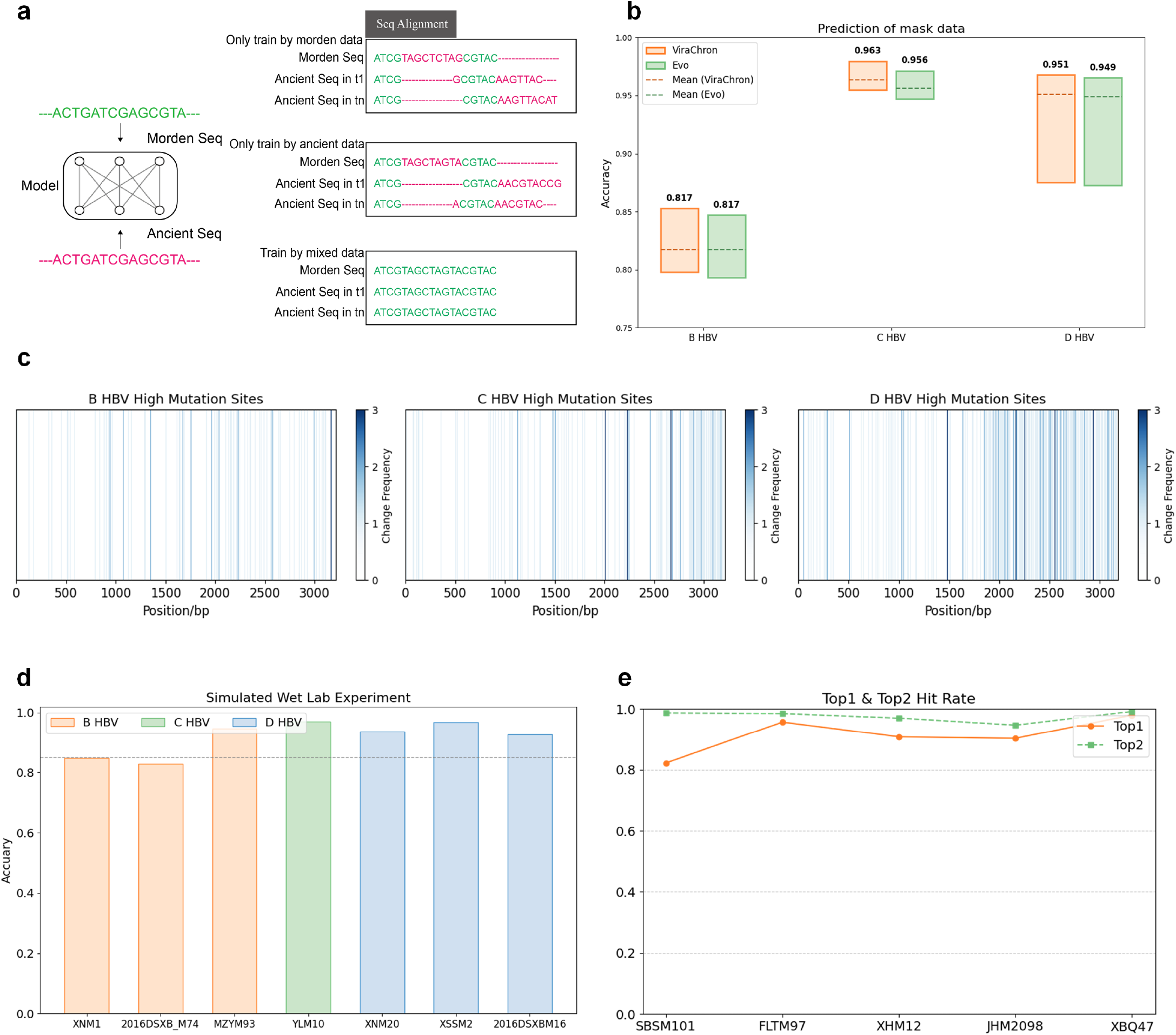
Testing the fundamental capabilities of ViraChron. (a) Fine-tuning approaches. Multiple sequence alignment (MSA) was performed on genome sequences completed by three different fine-tuning methods. Aligned genome sequences are shown in green, while misaligned genome sequences are shown in red.(b) Comparison of predictive performance of Evo and ViraChron. Both models were used to complete artificially masked sequences, with orange representing ViraChron and green representing Evo. The mean values shown here are weighted averages. (c) Statistical analysis of mutation sites.(d) Simulated wet-lab experiments. The model was used to impute missing genomic data from uncaptured sequences, and the quality of the imputed sequences was evaluated. (e) Nucleotide hit rate statistics for imputed sequences at corresponding BAM file positions.

### HBV evolutionary patterns

Although HBV evolution is functional rather than linear and sequential, it still exhibits temporal dependency[16]. We developed and used a multi-scale temporal encoding technique to learn sequence evolution patterns across different timescales (Methods). This encoding strategy is analogous to the positional encoding in traditional Transformer models[17], where discrete time information is transformed into temporally correlated tensors through multi-scale encoding.

To assess the impact of temporal information on model performance, we trained both Evo[7] and ViraChron for 500 epochs. This initial evaluation experiment did not account for the impact of genetic recombination, and variations in training data may substantially affect the model’s predictive performance. In addition, we also evaluated the ability of two models to capture the evolutionary dynamics of HBV. The results demonstrated that the time-augmented model outperformed the original model in overall accuracy (Figure 3b), achieving a higher average accuracy of 0.9173.

Stable sites in the HBV sequence primarily reflect fundamental genetic features of the genome, while highly variable sites reveal its evolutionary patterns. Our statistical analysis of common mutation sites across various HBV datasets (Figure 3c) revealed that only accurate predictions at these highly variable sites can effectively validate the model’s ability to reveal the genome evolution pattern.

We first assessed the model’s generalization ability. Based on the newly uncovered HBV sequences mentioned earlier, we predicted the uncaptured genome fragments and compared the predictions with the corresponding complete high-quality sequences (Figure 3d). The results showed that the model achieved a high prediction accuracy, ranging from 0.829 to 0.970, demonstrating a strong ability to generalize. To assess the model’s applicability, we used it to impute missing regions in 65 existing HBV sequences. Representative sequences were selected for comparison with their corresponding BAM files (Figure 3e). The results showed that the top2 accuracy ranged from 0.987 to 0.992, while the top1 accuracy ranged from 0.787 to 0.980. Specifically, we compared the imputed DA45 (Bayanbulag site) sequence with the high-quality AT19 (Bayanbulag site) data to assess the completion of missing regions in DA45 (DA45 and AT19 sequences are from the same individual). The imputed DA45 sequence showed only two discrepancies compared to AT19, demonstrating good imputation performance.

Ancient HBV genomes often lack a large amount of genetic information. We evaluated how the model performs in filling these gaps. Sequences from ancient HBV genotypes A, B, C, and D were completed, and the completed segments were extracted for a sequence similarity search to verify whether the model incorporated genomic patterns from other species (Methods). We then performed sequence identity analysis on the sequences before and after imputation and compared the genotypes corresponding to the HBV data before and after imputation (Figure S2). The results showed that the data before and after imputation belonged to the same genotype, with 81% sequences showing correct recognition in subgenotype classification.

To further validate the accuracy of the imputation, we conducted phylogenetic analysis to explore the evolutionary relationships among HBV sequences. We quantified the differences in the topological structure of different phylogenetic trees using the weighted Robinson-Foulds distance and constructed an MCC tree using BEAST software with the same dataset (Figure 4 and Figure S3 and RF distance table). By comparing the topological structure differences between trees generated from two independent runs as a control experiment, the results indicated that the topological structure difference between the phylogenetic trees before and after imputation (0.14) was greater than the difference observed between two parallel runs (0.107). However, compared to the positive control experiment, the difference introduced by the imputation was relatively small (0.033), and we consider this difference to be within a reasonable range since it does not significantly impact the overall phylogenetic analysis.

**Fig. 4.**
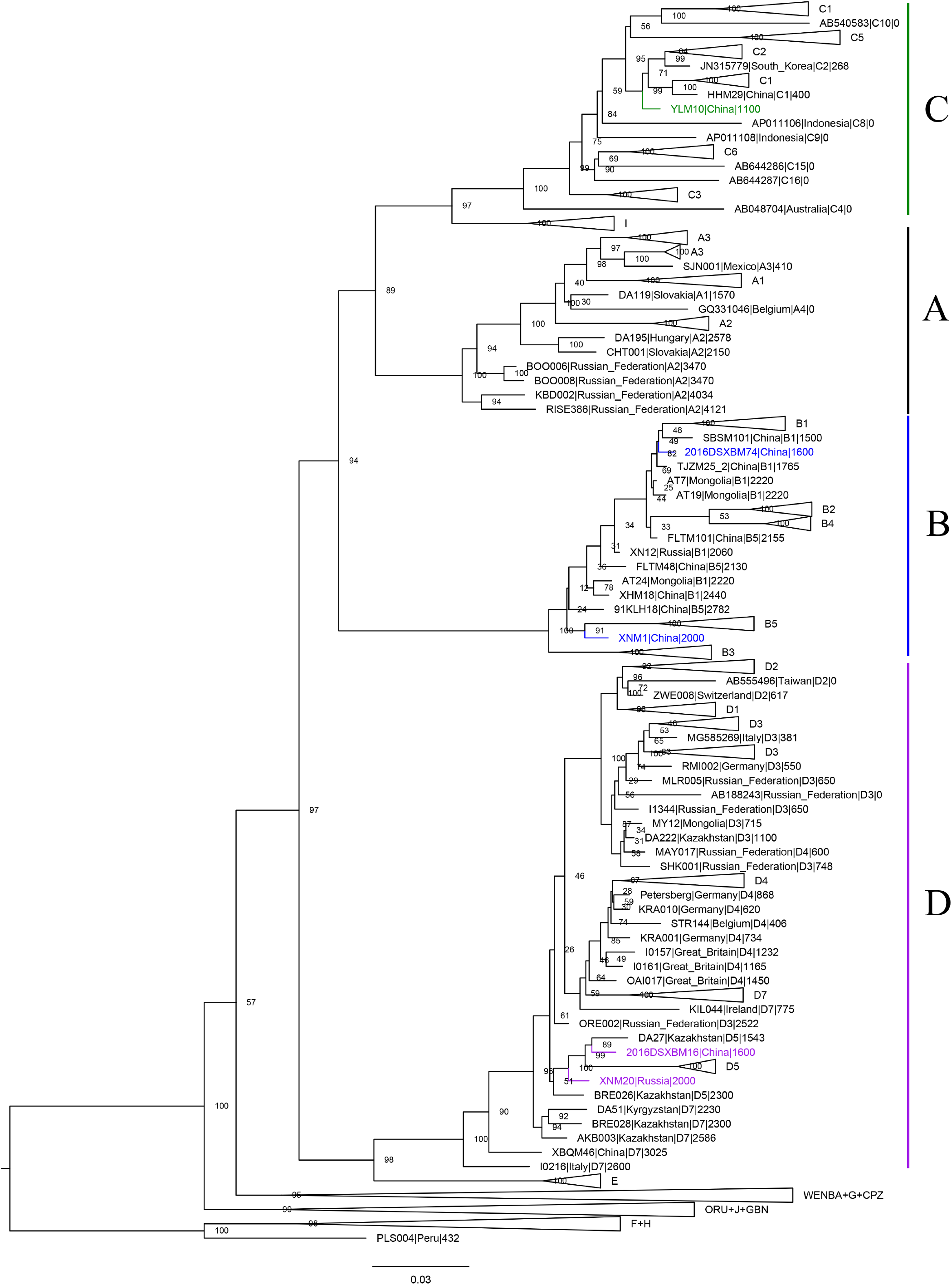
Maximum likelihood phylogenetic tree of modern and ancient HBV. The dataset used for phylogenetic tree construction included ancient sequences with coverage greater than 90%, which had been completed using our sequence-based foundation model. The lineages in which our ancient sequences are located are highlighted with different colors. The branches and names of the ancient sequences are also highlighted with the indicated color.

ViraChron results enabled the analysis of mutations and recombination in HBV evolution. Since mutation is a fundamental driving force of species evolution, tracing the changes in genetic material over time can estimate the genetic distance between individuals or populations, as well as the time since their most recent common ancestor. We calculated the base substitution rates for both the original and imputed datasets of HBV and performed Bayesian skyline analysis. The results showed that the base substitution rates remained consistent before and after imputation, and there were no significant differences in the demographic history of populations in the Bayesian skyline analysis (Figure 5). These findings suggest that sequence imputation did not affect the estimation of base substitution rates, thereby validating the reliability of this approach.

**Fig. 5.**
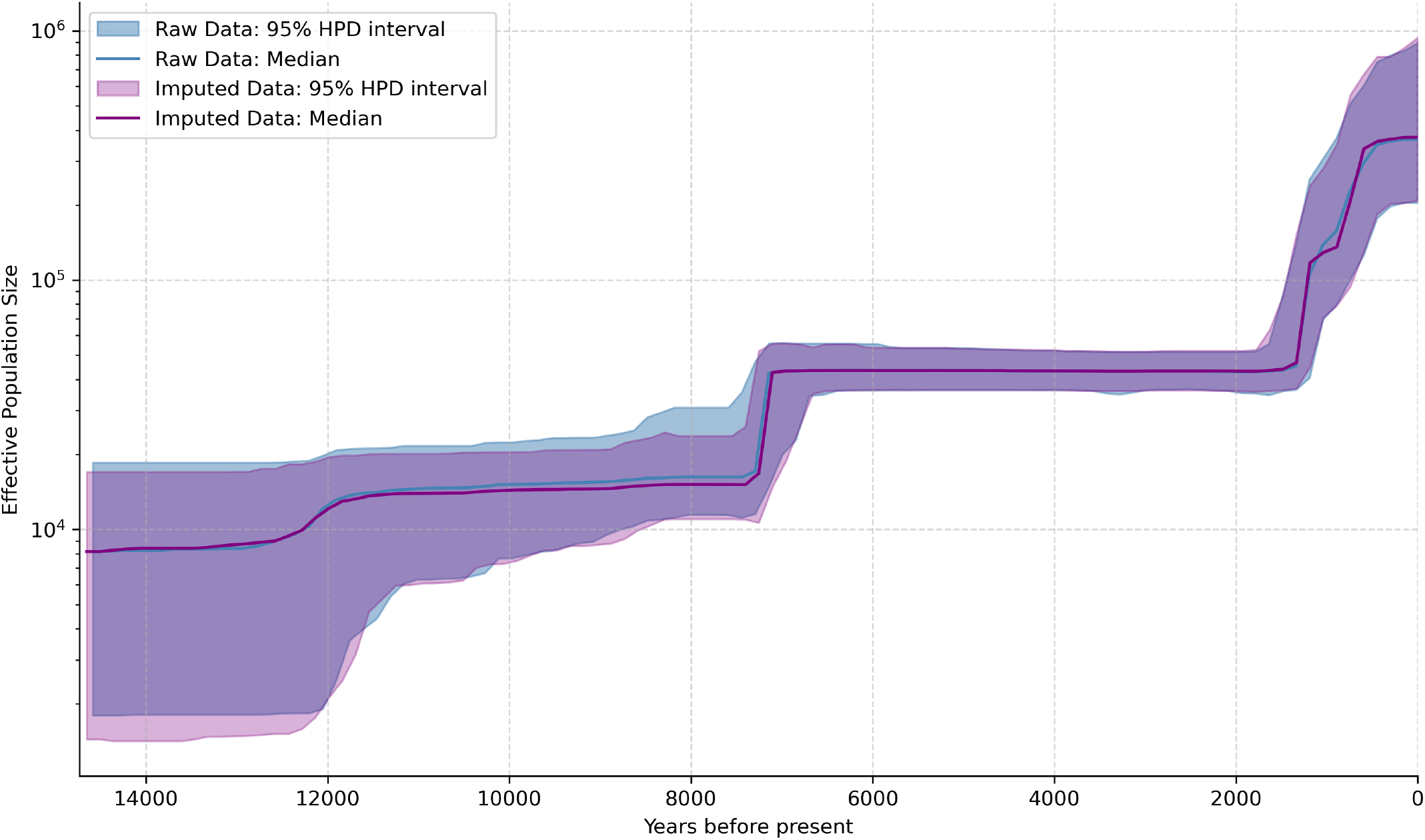
Bayesian skyline plot showing changes in the effective population size (Ne) of HBV over time, based on datasets before and after imputation, as estimated using the Bayesian Coalescent Skyline model implemented in BEAST[**?**]. Lines represent the median estimates for HBV, with shaded regions indicating the 95% highest posterior density (HPD) intervals.

### HBV mutation patterns

From a temporal perspective, the evolution of HBV is driven by adaptation to historical conditions and is based on past genetic patterns. To further analyze the temporal evolutionary pattern of this virus, we selected the oldest data available for each HBV genotype as a reference. The number of mutations in the imputed data at each time point was then calculated (Figure 6a and Figure S4). The results indicated that the mutation rate (Methods) of HBV gradually decreased over time, which was consistent with previous studies. This is due to the fact that HBV exhibits a faster evolutionary rate in the short term, driven by a high mutation rate and selective pressure, whereas in the long term, the evolutionary rate slows down due to stability and adaptive equilibrium.

**Fig. 6.**
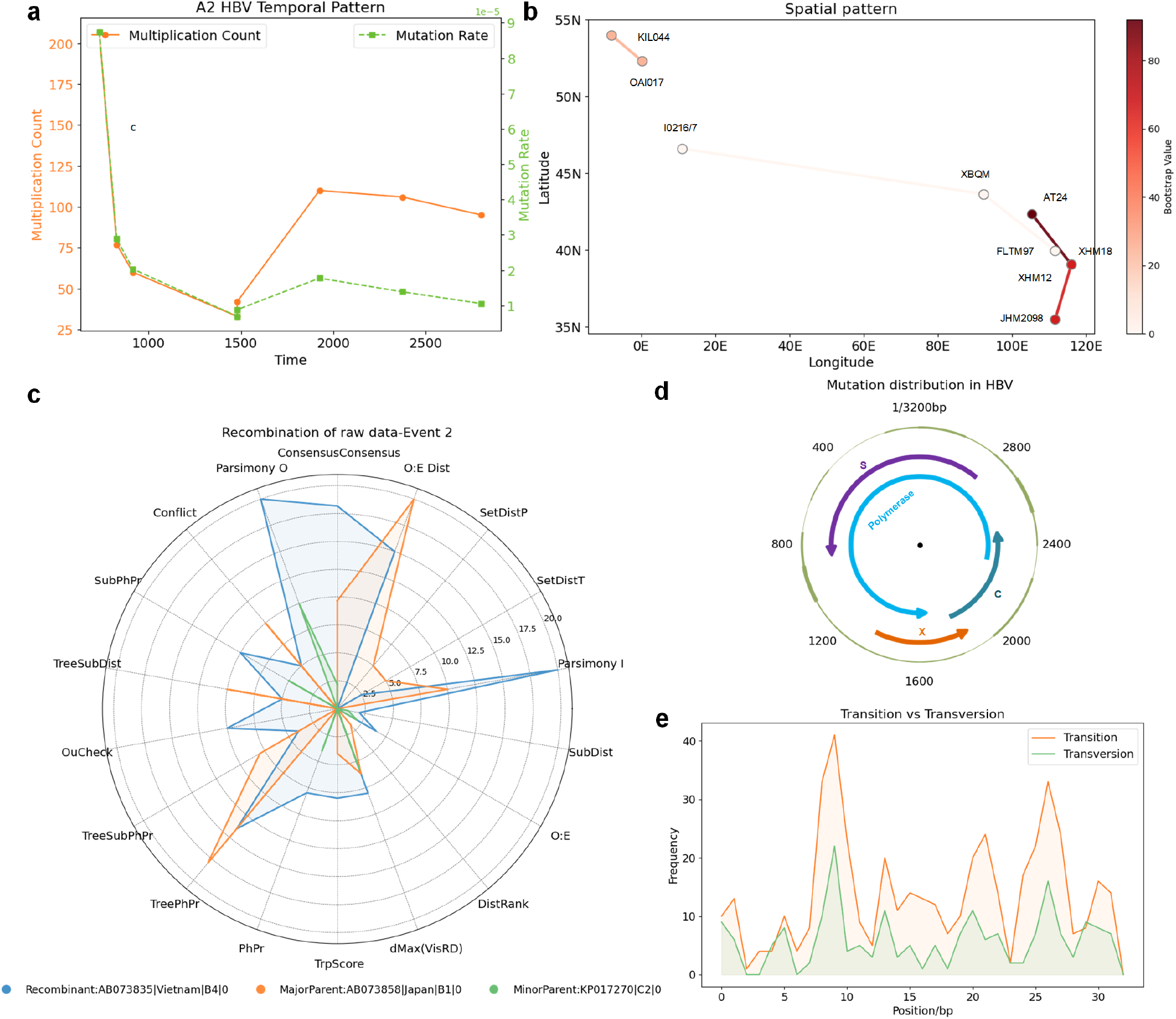
Evolutionary Patterns of HBV.(a) Temporal mutation patterns of HBV. Using the oldest data as a reference, the mutation count (orange) and mutation rate (green) for HBV at different time points were statistically analyzed. (b) Spatial mutation patterns of HBV. Circles represent individual sequences, and the colour of each connecting line denotes the bootstrap support of the corresponding branch in the maximum-likelihood phylogenetic tree. A credibility of 0 means the data does not belong to the primary branch but to a secondary branch. (c) Statistical analysis of mutation site information. The outermost circle (green) represents the HBV genome, and the thickness of the arcs indicates the number of mutation sites in each region.(d) Statistics on transitions and transversions.(e) Recombination information of the raw data-Event 2.

Additionally, we observed that some datasets exhibited significantly higher mutation rates, which may reflect the impact of large-scale human migration or environmental changes on HBV evolution during specific historical periods. From a global geographical perspective, HBV evolution involves transmission between different regions. When constructing a phylogenetic tree, genetic information loss may lead to incorrect classification of sequences with similar genomic patterns. We compared the branching changes in the phylogenetic tree before and after sequence imputation, and identified sequences with significantly reduced genetic distances after imputation. The results showed that the model can identify similar genomic patterns in the known portions of these sequences during the imputation process and generate corresponding imputed sequences (Figure 6b). Notably, the data from AT24 (Bayanbulag site) and XHM18 (Xihe site) showed high correlation after imputation (bootstrap support = 92%), and AT24 exhibited abnormal substitution rates in the temporal evolutionary pattern analysis. This is consistent with previous research findings[18]. The SBS site with individual AT24 is located in Mongolia and the Xihe site where individual XHM18 was found is located in northern China, making the two geographically distant. However, previous studies have proven that the individuals found at the SBS site were Han soldiers from northern China who were sent to guard the border[18]. Therefore, it is possible that the AT24 individuals became infected with HBV before their deployment, which could explain the similar evolutionary patterns observed between these two geographically distant sequences.

### HBV recombination patterns

Recombination also plays a crucial role in the genetic diversity and adaptive evolution of HBV[19, 20]. Specifically, virus strains from different genotypes or subgenotypes can exchange genetic material through recombination, leading to the formation of new viral variants that are better able to adapt to the host environment and immune pressure. To evaluate the impact of sequence imputation on analysis of recombination events, we performed recombination analysis using both the raw and imputed datasets. The results showed that the recombination events detected in both datasets occurred at similar times, including several widely validated recombination events: two recombination events in genotype B and a recombination event between genotype A, D and the WENBA lineage. This further supports that sequence imputation had minimal impact on recombination detection and did not introduce significant systematic bias. Additionally, the analysis of the imputed dataset revealed an extra recombination event within genotype D and two extra recombination events involving genotypes G and C (Figure 6c and see recombination_info-FINAL.xlsx). For one recombination event, four out of six detection methods yielded P-values between 0.01 and 0.05, indicating moderate support. Recovery of a high-coverage genome through target enrichment using hybridization capture typically requires approximately 80-90 hours. In comparison, the model developed in this study completes the reconstruction of a sequence with 50% coverage in 5.4 minutes, corresponding to an average of 0.2 seconds per nucleotide, achieving an 889-to 1000-times increase in speed relative to traditional approaches.

### HBV mutation hotspots

To further explore the natural evolutionary patterns of HBV. We used the 7804 modern HBV sequences used initially to train our model and compared them with ancient HBV data. The results successfully identified known HBV mutation hotspots, for example, in genotype B HBV, all ancient sequences exhibited an isoleucine substitution at position 204 in the P region (rtM204I). In genotype A HBV, we observed a fixed threonine-to-alanine mutation at position 1850 in the pre-C region of the ancient sequences. Additionally, we systematically analyzed regions with higher mutation frequencies in ancient HBV (Figure 6d) and found that all genotypes HBV exhibited a high mutation rate at the end of the P gene (approximately 850–1050 bp)(Figure S5). This phenomenon may be influenced by specific biological mechanisms or historical factors.

Further analysis revealed a regular distribution of transitions and transversions at specific positions in the HBV genome (Figure 6e), which not only demonstrates the prevalence of HBV mutations but also reveals that its mutational patterns exhibit site-specificity and particular changes, rather than a completely random and uniform distribution across the genome.

Overall, our model effectively captures the evolutionary dynamics of HBV. Despite the challenges of incomplete data, the imputed sequences preserved key evolutionary features, only introducing minor distortion to phylogenetic relationships and recombination patterns.

## Discussion

In this study, we developed and tested an innovative temporal foundation model specifically designed for the task of ancient sequence imputation. A unique feature of this model is the incorporation of temporal signals for the first time, an innovation that significantly enhanced its performance. Through in-depth analysis of ancient HBV genomes before and after imputation, we validated the robustness of the model on highquality data, providing more comprehensive and reliable data support for understanding HBV evolution. The seven newly published ancient HBV sequences from this study, imputed with our model, offered an unprecedented opportunity for detailed evolutionary investigation.

Capitalizing on this high-quality dataset, we compared the imputed oldest sequences with all HBV sequences in the NCBI database to identify conserved mutations. Conserved mutations are those present only in the oldest sequences but replaced in all other data. In genotype A HBV, we identified multiple conserved mutations and analyzed their potential impact on protein function (Figure S7). Notably, we detected a premature stop codon in the P gene, a phenomenon not observed in other genomes. The hepatitis B virus polymerase protein is one of the core proteins of the virus, playing a critical role in viral genome replication. If a nonsense mutation occurs in the gene encoding the polymerase protein, preventing its normal expression, it would hinder viral reverse transcription and DNA synthesis, disrupting genome replication and virion assembly, ultimately affecting the viral life cycle. Given the profound biological implications of such a finding, we critically assessed the authenticity of the mutation. A closer examination revealed that the sequencing reads supporting this mutation did not exhibit the typical damage patterns of ancient DNA. This rigorous validation process allows us to distinguish true viral signals from off-target noise originating from the metagenomic background, thereby ensuring the integrity of the biological conclusions reported in this study.

### HBV transmission with wars leading to the coexistence of genotypes B and D

The newly published sequences belong to genotypes B, C, and D, with the coexistence of genotypes B and D first observed at the Tiantiajie and Derestuj sites. Previous studies indicated that genotype B infections were present at the Tiantiajie site, while genotype D infections were primarily found in nomadic populations[10]. Our findings align well with the historical context of the Tiantiajie site as the capital of the Northern Wei Dynasty, Pingcheng. The rulers at that time were the Xianbei people, who were proven to carry genotype D HBV. The rulers recruited a large number of foreign laborers for capital construction, creating favorable conditions for HBV exchange and transmission[21]. Similarly, the Derestuj site, associated with the Xiongnu culture, also provided favorable conditions for HBV exchange and transmission due to the wars between the Xiongnu and the Han, leading to the coexistence of genotypes B and D[22]. In contrast to the widespread mixing observed for genotypes B and D, the distribution of ancient genotype C appears to have been more geographically restricted. While mainly found in Northeast China and Korea, the genotype C identified in this study was from noble individuals buried in Yelüyuzhi. Previous studies have linked ancient genotype C samples to populations such as the Jurchen descendants. Crucially, despite the known presence of Xianbei-related populations in regions where genotype C circulated, we found no evidence of genotype C at the major Xianbei center of Tiantaijie (Pingcheng). This absence suggests that the large-scale migration of Xianbei populations to establish their capital was not a significant vector for the transmission of genotype C, reinforcing the hypothesis of its limited geographical spread during that period.

### ViraChron reveals interactions among the local population that facilitated HBV evolution

We performed sequence imputation on data with coverage greater than 90% and conducted phylogenetic analysis based on the imputed data. Additionally, to comprehensively evaluate the evolutionary relationships of ancient HBV, we constructed a phylogenetic tree using both raw ancient sequences with coverage greater than 50% and modern sequences. The analysis revealed that the genotype B sample 2016DSXB_M74 (Tiantaijie site) clustered with other individuals from the same site on a small branch of the phylogenetic tree, suggesting a common source of genotype B HBV infection at this site. This finding reflects close interactions among the local population at that time, facilitating the spread and evolution of HBV within the region. The genotype B sequence XNM1 from the Derestuj site did not cluster with sequence XNM12 from the same site, indicating higher HBV diversity in this region, potentially due to a more complex history of population mixing. Furthermore, individuals from the Meiziyuan site clustered with those from the FLT site on the phylogenetic tree, supporting the possibility of HBV transmission between different sites.

### Filling a gap in the genomic diversity of genotype C

In the analysis of genotype C, the individual identified in this study dates back to 1100 AD, filling a gap in the genomic diversity of genotype C during this period. This sequence is located at the root of branches C2 and C1, representing diversity prior to the divergence of C2 and C1, and provides important insights into the early evolution of genotype C HBV. For genotype D, ancient samples XSSM2 (Shihuyao site), 2016DSXBM16 (Tiantaijie site), and XNM20 (Derestuj site) were located within a large branch that also includes modern D5 sequences and ancient samples from the Siberian Plain, Mongolian Plateau, and Tianshan region. This result suggests a high degree of similarity among genotype D HBV infections in these regions, likely reflecting frequent population migrations and cultural exchanges throughout history.

Overall, through in-depth analysis of ancient HBV genomes before and after imputation, we validated the robustness of ViraChron on high-quality data, providing more comprehensive and reliable data-driven support for understanding HBV evolution. Our innovative and efficient approach uncovered seven newly published ancient HBV sequences recovered from sites in China and Russia. These sequences belong to genotypes B, C, and D, with the coexistence of genotypes B and C observed at the Tiantaijie and Derestuj sites. This discovery provides new evidence for the spread and evolution of HBV. Nevertheless, with the emergence of more high-quality ancient DNA data, we anticipate that the performance of this model will further improve, thereby advancing our understanding of viral evolution in humans.

## Methods

### DNA extraction and library preparation

Ancient DNA work was carried out in dedicated cleanroom laboratory facilities at the ancient DNA laboratories of Jilin University in Changchun. Teeth were drilled and powder was collected. A total of 50mg of tooth or pars petrosa powder was used for extraction following an established protocol (https://dx.doi.org/10.17504/protocols.io.bqebmtan). The extracted DNA was transformed to generate double-stranded genetic libraries. Genetic libraries were indexed and amplified before shotgun sequencing. In addition, negative controls were taken along with initial library preparation. These libraries were shotgun sequenced on an Illumina HiSeq X10 or HiSeq 4000 instrument using 2 × 150-base-pair (bp) chemistry.

### Screening and enrichment experiment

Before performing alignment and taxonomic binning of the obtained reads with MALT (v.0.5.3), each sample was mapped to the human reference genome (hs37d5) using EAGER1[23, 24]. The reads that did not map to human sequences were extracted from the bam files using samtools (v.1.3) (samtools view -f 4)[26]. Finally, we used bedtools bamtofastq (v.2.25.0) to convert the bam file into a fastq file26. These non-human reads were taxonomically assigned by MALT with the reference datasets containing known modern HBV diversity as well as other orthohepadnaviruses. All runs used ‘semi-global’ alignment and a minimum percent identity of 90. For samples that had reads mapped to HBV in the MALT analysis, we used reference sequences comprising multiple HBV genotypes for comparison using bwa, so as to once again count the reads belonging to HBV in the sample metagenome. To identify the genotype of these individuals, we performed a competitive mapping with a combined reference using the EAGER pipeline. Finally, we counted the reads mapping to each sequence to determine which is the most likely genotype for each of the samples.

After screening, those libraries identified as positive for HBV were enriched for HBV DNA using an in-solution target enrichment of HBV following the strategy used in previous ancient HBV work[27]. The HBV probes were designed by iGeneTech Co. Ltd (Kit name: AI-HBV-Cap Enrichment Kit, article number: AIHBC), and the experiment was conducted following the manufacturer’s instructions.

### HBV genome reconstruction

After determining the genotype of each individual, we chose a reference and repeated the mapping steps described above. SNP and INDEL calling was carried out with Genome Analysis Toolkit (GATK) UnifiedGenotyper version 3.5 using a quality score of ≥30 and the “EMIT_ALL_SITES” output mode[28]. Then, consensus sequences were created using GenConS, which is available in the TOPAS package (major_allele_coverage 3, consensus_ratio 0.9, punishment_ratio 0.8) (https://github.com/subwaystation/TOPAS).

### Analyzing damage patterns

After determining the genotype of each individual, we chose a reference and repeated the mapping steps described above. To check for the presence of damage patterns characteristic of ancient DNA, i.e., the accumulation of C > T changes due to C deamination at the 5’ end of the fragments, we used mapDamage v.2.0.9-dirty with default parameters[29]. Except for individuals with only a few HBV reads in shotgun data and those for whom full-UDG treatment of the libraries was performed for the in-solution capture experiment, all other genomes showed damage patterns typical of ancient DNA in the reads mapping to the HBV genome.

### Sequence identity of the dataset before and after imputation

We used the imputation method developed in this study to recover all 65 published sequences as well as the newly identified 7 HBV sequences published here. To verify the accuracy of the imputation model, we conducted sequence identity analyses on ancient sequences before and after imputation. The reference sequences used in the analysis were modern HBV sequences from different sub-genotypes, and the same methods and parameters were applied in both analyses. For this we computed the number of insertions, deletions, and mismatches between modern and ancient sequences normalized by the total length of the sequence. Missing data in the ancient sequences were not included in the calculations.

### Phylogenetic analysis of the dataset before and after imputation

To validate whether the positioning of the imputed ancient samples on the phylogenetic tree shifts, we constructed maximum clade credibility (MCC) trees using two datasets. One dataset included sequences of ancient samples with imputed data together with modern HBV sequences, while the other dataset consisted of the pre-imputed sequences of the same ancient samples and the modern sequences. For time-calibrated phylogenetic analysis, we used radiocarbon dates of the ancient HBV genomes as calibration points in the BEAST analysis[30]. The molecular clock was calibrated using tip dates, where modern sequences were assigned a date of 0, and ancient sequences were assigned mid-range values based on 14C or archaeological dating. We employed a Gamma distribution for the site model, the GTR substitution model, and the relaxed log-normal molecular clock with coalescent exponential population priors. A uniform prior between 10-9 and 10-3 substitutions per site per year was applied for the mean clock rate, based on previous estimates. The total Markov chain length was set to 500 million. After the phylogenetic trees were generated, we used TreeAnnotator v2.6.2 to construct the MCC tree, discarding the first 10% as burn-in. All parameters achieved an effective sample size (ESS) greater than 200.

To quantify the topological differences between the phylogenetic trees constructed from imputed and non-imputed sequences, we used the Weighted Robinson-Foulds distance. Additionally, we ran the BEAST analysis with identical datasets to generate maximum clade credibility trees, and the topological differences between the trees produced by two separate runs were used as a positive control for comparing the topological differences.

### Bayesian skyline plot of the dataset before and after imputation

To analyze the dynamics of population size over time, a Bayesian Skyline Plot (BSP) was employed using BEAST2 with MCMC algorithms[30]. The prior was modified to a coalescent Bayesian skyline, while other parameters remained consistent with those used in the construction of the Maximum Clade Credibility (MCC) tree. The dataset used for this analysis was the same as that used for the MCC tree construction. The resulting log file was analyzed in Tracer to ensure Effective Sample Size (ESS) values were 200 or greater, and the BSP was subsequently visualized and extracted.

### Recombination analysis of the dataset before and after imputation

The recombination detection program version 5 (RDP5) was used to search for evidence of recombination within all newly published and previously published ancient sequences, a selection of 134 modern HBV sequences, and non-human primate sequences[31]. We performed two rounds of recombination detection analysis, where the newly published and previously published ancient samples were analyzed separately before and after sequence imputation. Seven recombination methods (RDP, GENECONV, BootScan, MaxChi, Chimaera, SiScan, and 3Seq) were used to detect the recombination event with default parameters. In this analysis, RDP5 constructed maximum likelihood trees for each recombination event separately, using different regions from the presumed major and minor parents in the recombinant. The authenticity of recombination events was confirmed by comparing the position of the recombinant in these two ML trees.

### Model Training

Our training dataset comes from NCBI, covering complete HBV genomes of genotypes A, B, C, and D. After data collection, we initially filtered the sequences based on the length of the ancient HBV genomes. Next, we performed whole-genome alignment using MAFFT and excluded sequences with large differences from the ancient HBV genomes, thus constructing the model training dataset. To prevent the features of ancient HBV from being overshadowed by modern data during training, we implemented a data balancing strategy for each HBV genotype. During each training iteration, 200 modern HBV sequences were randomly selected from the entire training set and input into the model, even though time features had already been introduced, to ensure the effective learning of ancient HBV features.

We adopted the evo-8k-1-base pretrained parameters and fine-tuned them using LoRA (Low-Rank Adaptation) for the MLP in the framework. The Deepspeed framework was used to support efficient training across 10 NVIDIA RTX 3090 GPUs. During model training, we split the HBV genomes into 500 bp intervals and trained with a batch size of 1. The learning rate was set to 0.00002, and we employed a warm-up mechanism to gradually increase the learning rate during the first 200 steps to stabilize the training process. Furthermore, after each training epoch, we processed the inf values and introduced a smoothed cross-entropy loss to mitigate gradient anomalies and improve the model’s prediction stability and generalization ability.

### Time Encoding

Each input genome sequence was represented as a nucleotide sequence of shape ℝ^*n*×1^, where *n* was the sequence length. Nucleotides (A, T, C, G) were first mapped into a 4096-dimensional latent vector using a tokenizer and a high-dimensional embedding layer, resulting in a representation of shape ℝ^*n*×4096^. To incorporate temporal context, we introduced a time variable *t*, which denoted the chronological information associated with each sequence, such as sampling time or evolutionary age. Due to the large temporal span across sequences, we applied logarithmic normalization to *t* to reduce numerical instability and satisfy evolutionary time dependency:

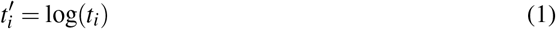

We then encoded the normalized time 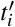 using an exponential decay sine/cosine transformation across multiple frequency scales:

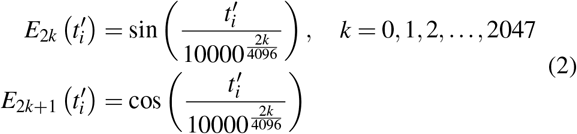

The resulting time encoding vector was denoted as ***E*** ∈ ℝ^1×4096^. To align with the nucleotide embeddings along the sequence length dimension, this vector was replicated *n* times, yielding a time encoding matrix ***E***′ ∈ ℝ^*n*×4096^.

To optimize the integration of temporal information while preventing it from overwhelming the core genomic representations, we controlled the contribution of the time encoding by applying a scaling factor. Specifically, we tested different weights (0.1, 0.02, 0.01) for ***E***′ during fusion with the latent representation to balance evolutionary context and nucleotide identity.

### Mutation rate

The mutation rate of the genome was calculated using the earliest sample of each genotype as a reference.

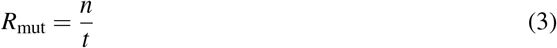

Here, *n* represented the number of genomic differences relative to the reference, and *t* denoted the evolutionary time.

### Gene Prediction and Malignant Mutation Analysis

We generated the corresponding four genes based on the HBV genome information discussed earlier and performed alignments using BLAST(2.12.0+)[32] to determine whether the imputed sequences contained genes from other species. We analyzed the distribution of the four genes in the complete HBV genome and conducted mutation sensitivity analysis using a sliding window strategy. Specifically, we selected a window every 100bp and randomly introduced in-silico mutations at 5bp, 10bp, or 50bp intervals within this window. Each mutated nucleotide was randomly replaced with A, T, C, or G with equal probability to simulate sequence noise and assess BLAST’s sensitivity to noisy genes.

To minimize the impact of the full HBV genome on BLAST homology analysis, we only extracted the imputed regions for alignment. Specifically, we selected continuous segments of imputed sequences longer than 20bp and used them as query sequences for BLAST homology searches to detect the source of the imputed sequences.

### Model Inference

During inference, we selected a 200 bp upstream sequence fragment of the missing site as input and injected multi-scale temporal information. For the probability tensor generated by the model, we chose the nucleotide corresponding to the maximum probability as the base for the imputed site and used it as the new context for iterative prediction. For HBV’s circular DNA structure, we performed dual predictions for incomplete sites with insufficient upstream sequences to improve prediction accuracy. During inference, we set the temperature parameter to 0.5 to enhance the model’s adaptability to HBV evolutionary patterns while maintaining a certain level of prediction diversity. All inference tasks were completed on an NVIDIA RTX 3090 GPU.

## Reporting summary

Further information on the research design is available in the Nature Research Reporting Summary linked to this article.

## Supporting information

RF distance table

recombination_info-FINAL.xlsx

## Code Avaliability

All source codes used in our experiments have been deposited at https://github.com/qiji24-jlu/ViraChron

## Acknowledgements

This work was supported by the National Science and Technology Major Project (No. 2021ZD0112500) and the National Natural Science Foundation of China (No.42372017 (Y.Q.), No. 62172187, No62372494, and No. 62372209); the Science and Technology Planning Project of Guangdong Province No. 20200708112YY and No. 2020A0505100018, and Guangdong Universities’ Innovation Team Project No. 2021KCXTD015 and Guangdong Key Disciplines Project No. 2021ZDJS138 and 2022ZDJS139.

