## Supplementary material for "Temporal foundation model unveils ancient Hepatitis B virus evolution": figs1-s7 tables1

**Table S1.** Summary of information for the individuals from which ancient HBV genomes were recovered in this study

| | XNM1 | MZYM93 | 2016DSXB_M74 | YLM10 | XSSM2 | 2016DSXBM16 | XNM20 |
|---|---|---|---|---|---|---|---|
| **Country** | Russia | China | China | China | China | China | Russia |
| **Site Name** | Derestuj | MeiZiYuan | TianTaiJie | Yelvyuzhi | ShiHuYao | TianTaiJie | Derestuj |
| **Lat** | 50.6 | 32.8 | 37.88 | 43.86 | 44.33 | 37.88 | 50.6 |
| **Lon** | 105.62 | 110.8 | 111.07 | 120.06 | 86.03 | 111.07 | 105.12 |
| **Period** | Early Iron Age | Late bronze age | Historical Period | Historical Period | Historical Period | Historical Period | Early Iron Age |
| **Skeletal Element** | tooth | tooth | tooth | tooth | tooth | tooth | tooth |
| **Library** | double-stranded | double-stranded | double-stranded | double-stranded | double-stranded | double-stranded | double-stranded |
| **UDG treatment** | NON-UDG | NON-UDG | NON-UDG | NON-UDG | NON-UDG | NON-UDG | NON-UDG |
| **Trac** | 7627953 | 5917436 | 6746645 | 7311391 | 7737077 | 5258693 | 14083605 |
| **RmPacR** | 11884 | 806 | 15658 | 2059 | 314 | 8035 | 9881 |
| **Mcac** | 304.1082 | 13.3621 | 495.2286 | 72.1997 | 11.5971 | 221.1025 | 266.204 |
| **1 ×** | 99.44 | 98.38 | 100 | 99.97 | 78.66 | 98.18 | 97.86 |
| **3 ×** | 97.85 | 81.9 | 100 | 98.16 | 71.81 | 97.74 | 97.52 |
| **Genotype** | B | B | B | C | D | D | D |

Trac: Total reads after capture.
RmPacR: Reads mapping to plague after capture after RMDup.
Mcac: Mean Coverage after capture

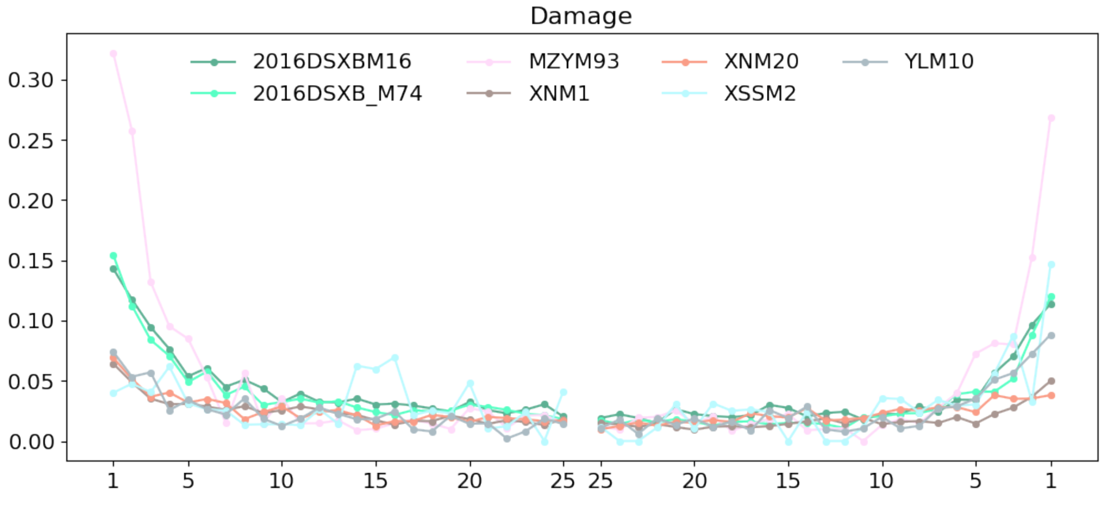


**Fig. S1.** Damage pattern of all the HBV positive samples to HBV.

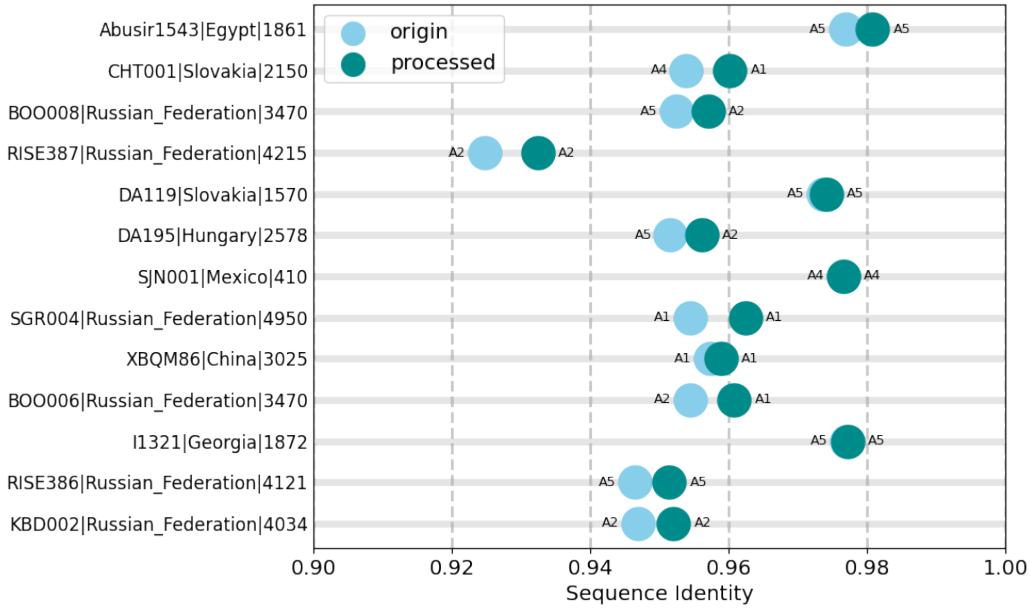


(a): Sequence identity analysis of genotypes A

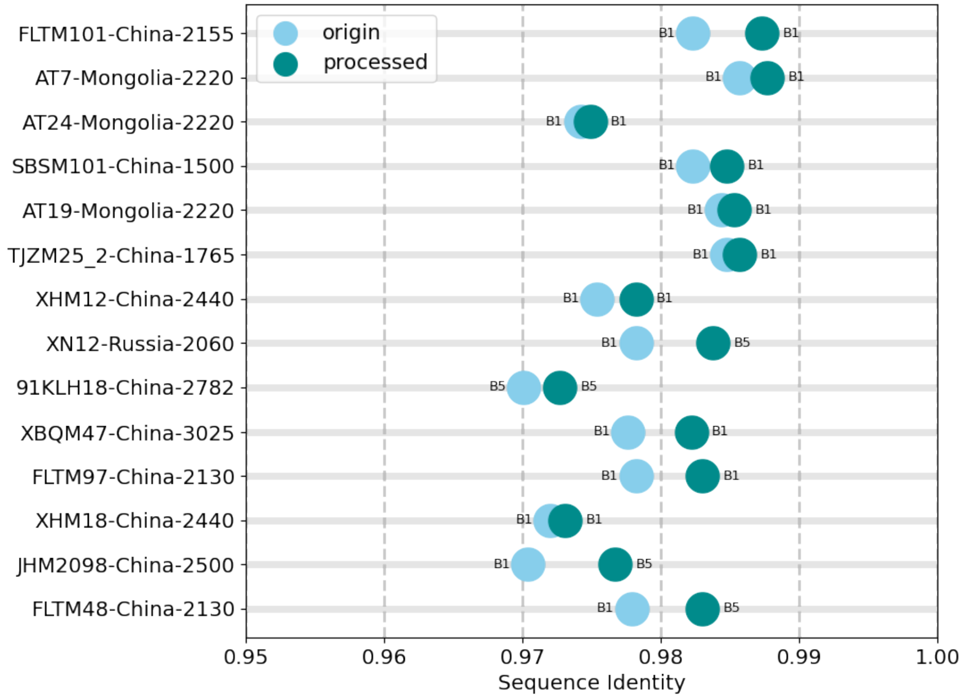


(b): Sequence identity analysis of genotypes B

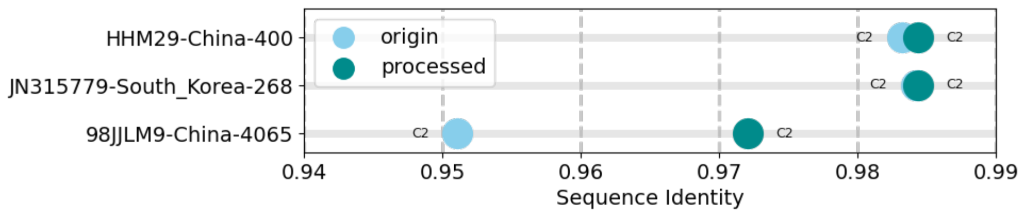


(c): Sequence identity analysis of genotypes C

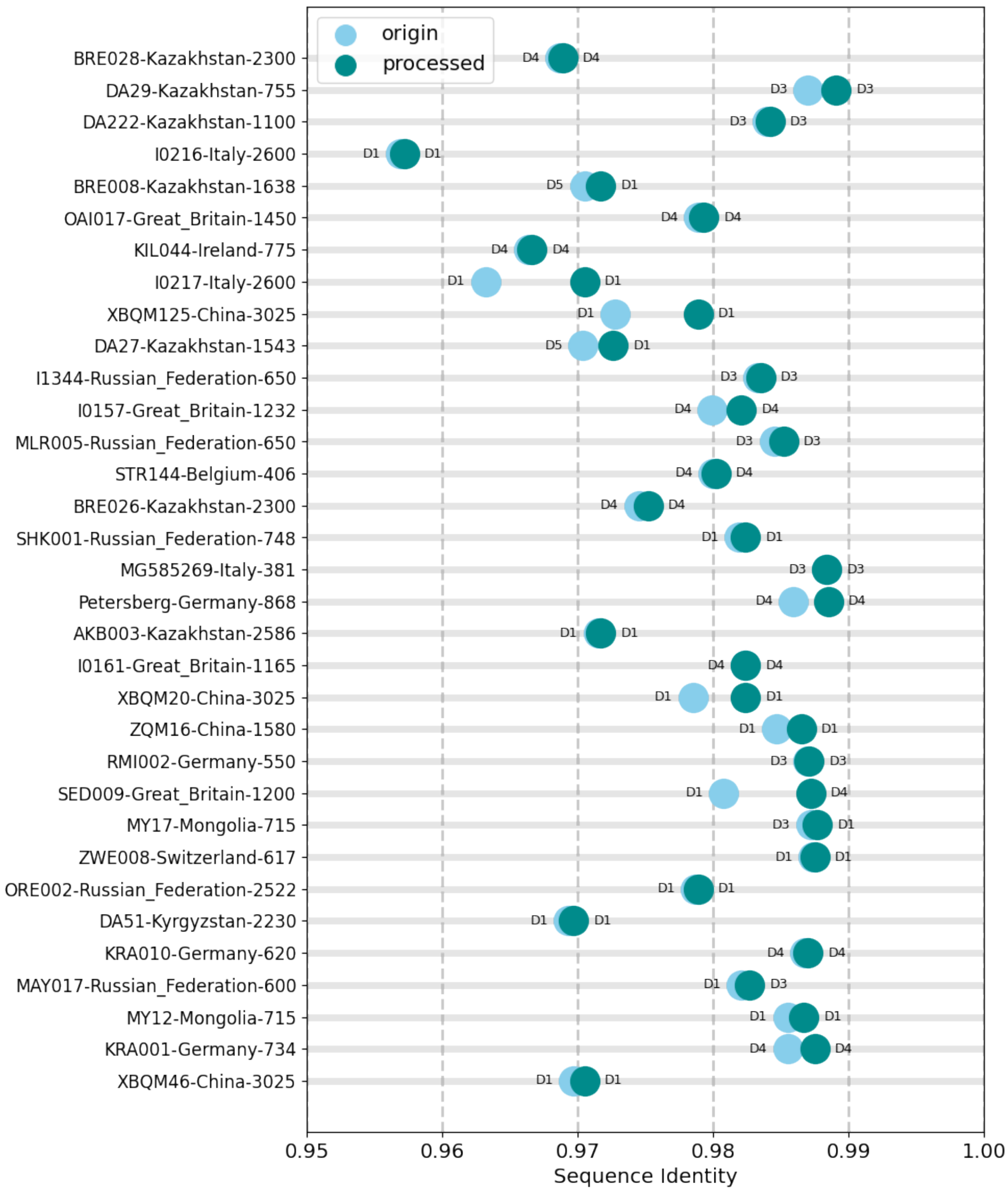


(d): Sequence identity analysis of genotypes D

**Fig. S2.** Sequence identity analysis of genotypes A, B, C and D.

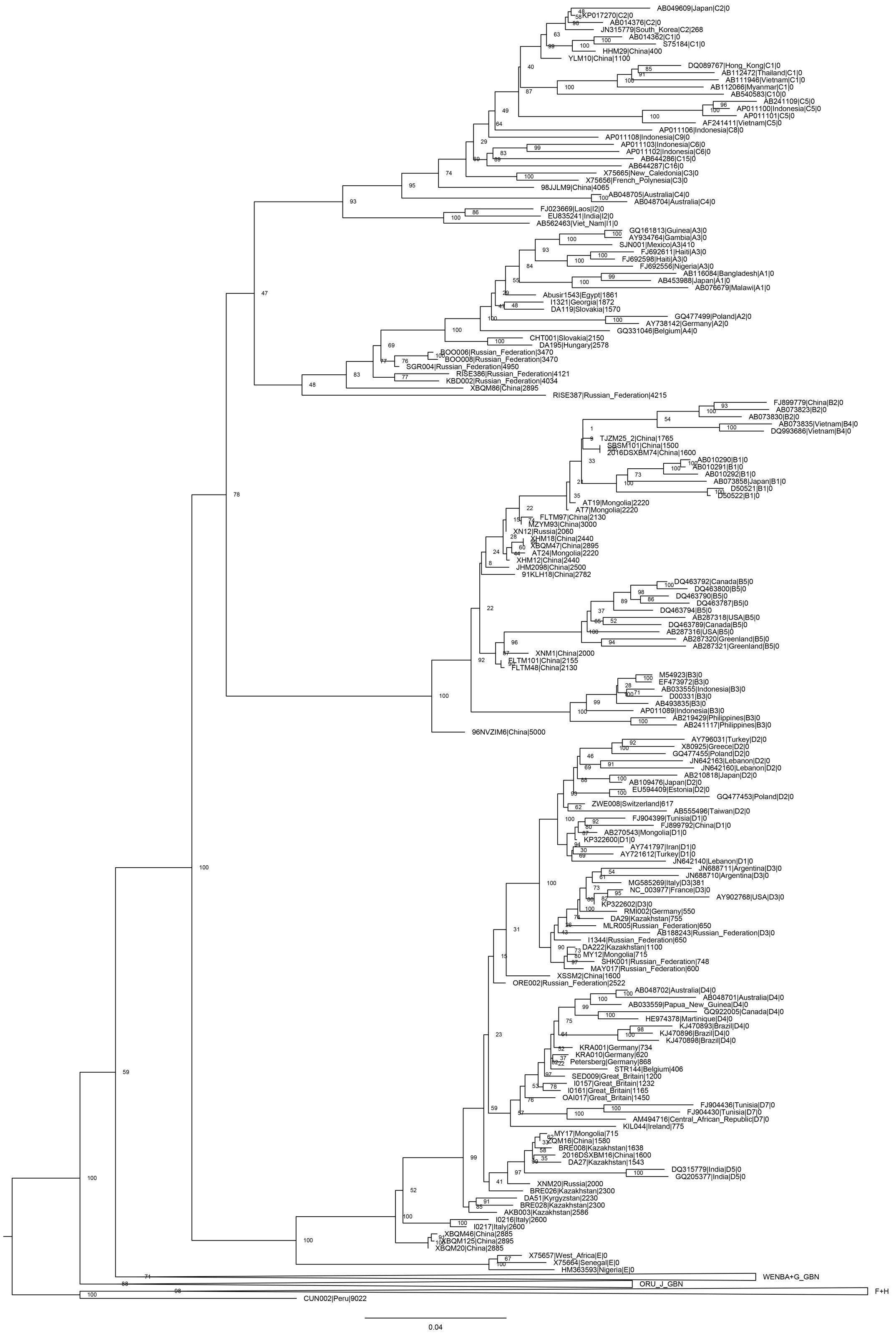

(a): Maximum likelihood tree based on the 50% coverage dataset

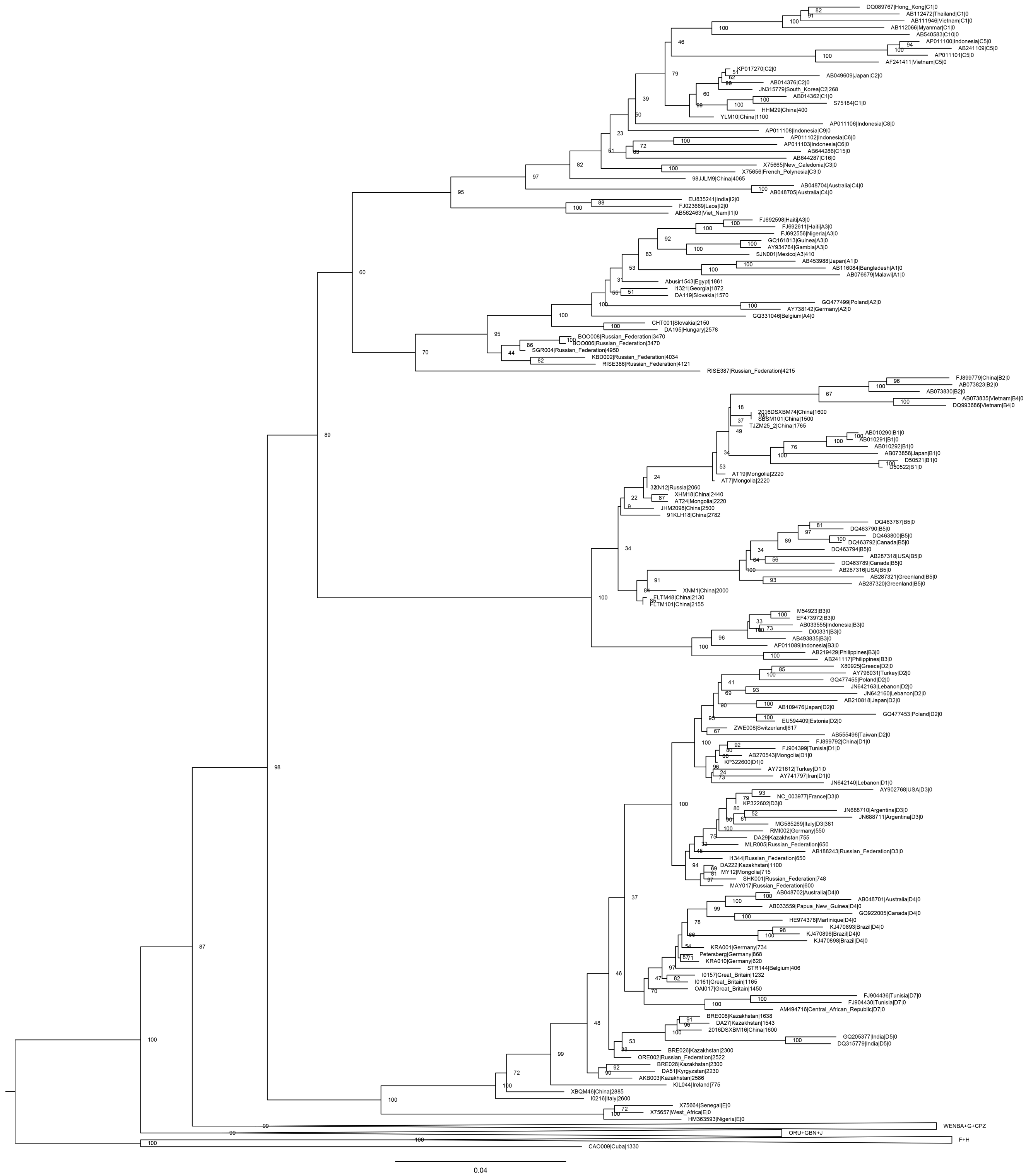

(b): Maximum likelihood tree based on the 80% coverage dataset

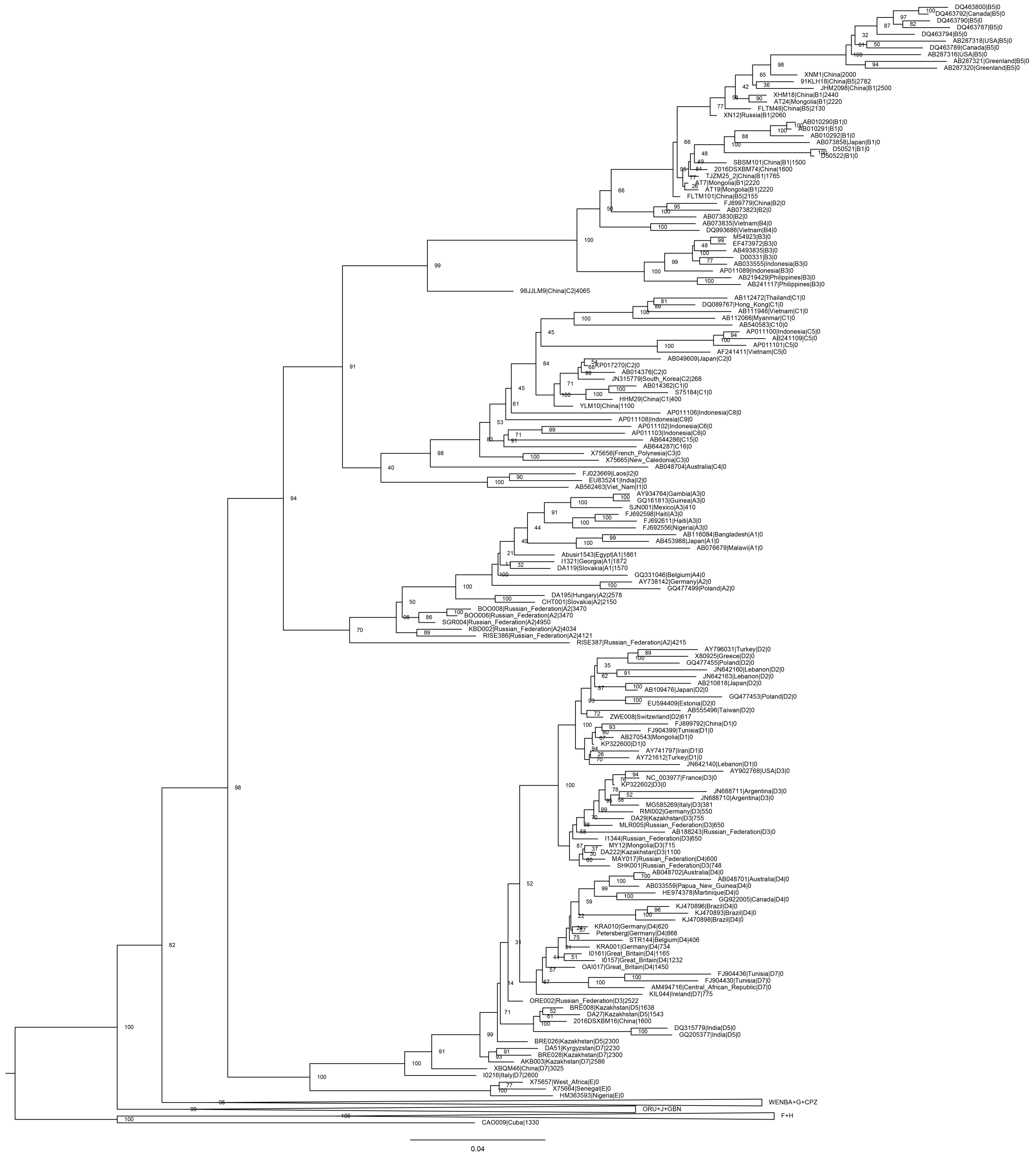


(c): Maximum likelihood tree based on the imputed dataset with minimum 80% coverage

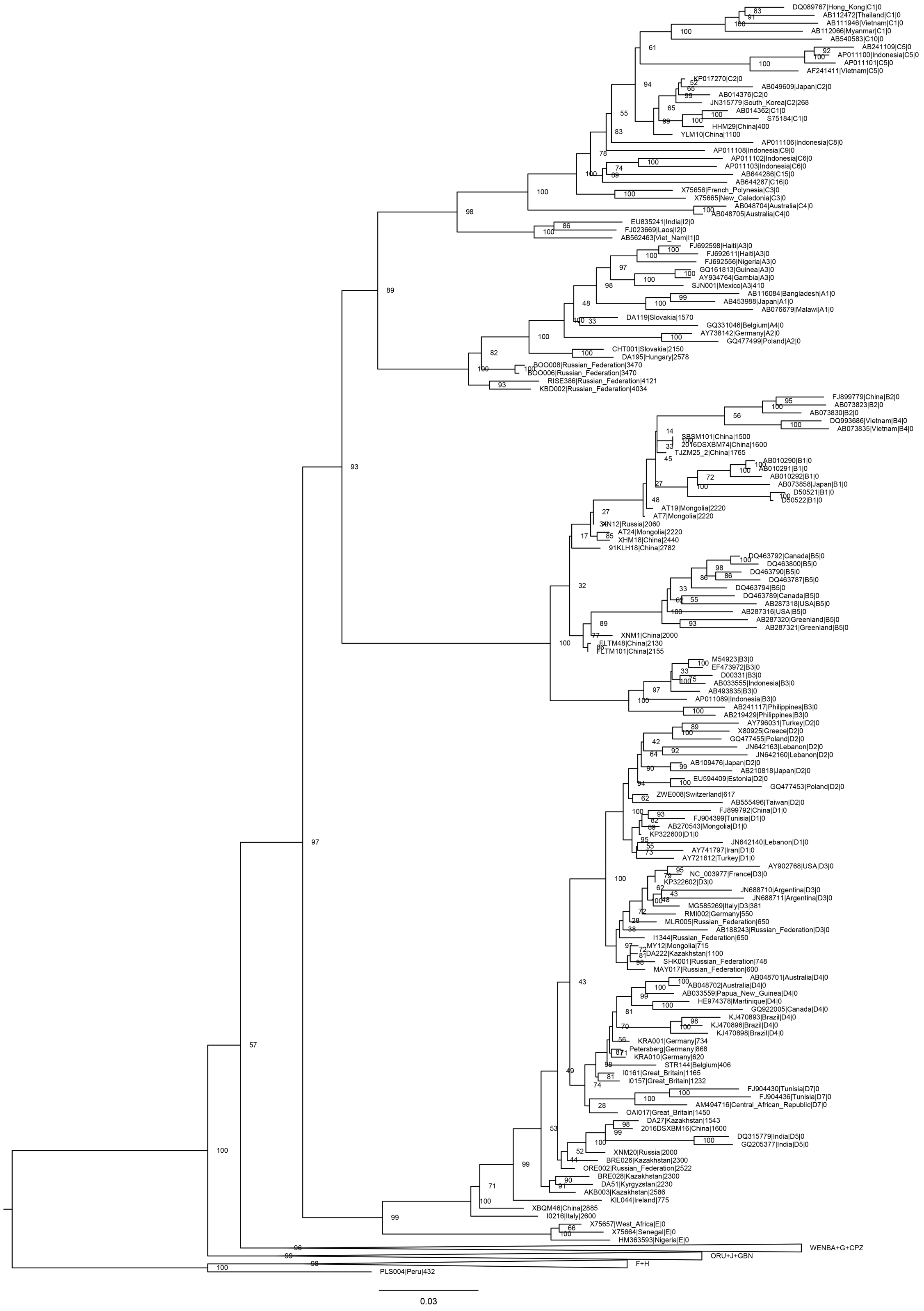


(d): Maximum likelihood tree based on the 90% coverage dataset

**Fig. S3.** Maximum likelihood phylogenetic tree of different datasets. (a) The dataset used for phylogenetic tree construction included the raw ancient sequences with coverage greater than 50%, (b) The dataset used for phylogenetic tree construction included the raw ancient sequences with coverage greater than 80%, (c) The dataset used for phylogenetic tree construction included ancient sequences with coverage greater than 80%, which had been imputed using our sequence-based large model, (d) The dataset used for phylogenetic tree construction included the raw ancient sequences with coverage greater than 90%.

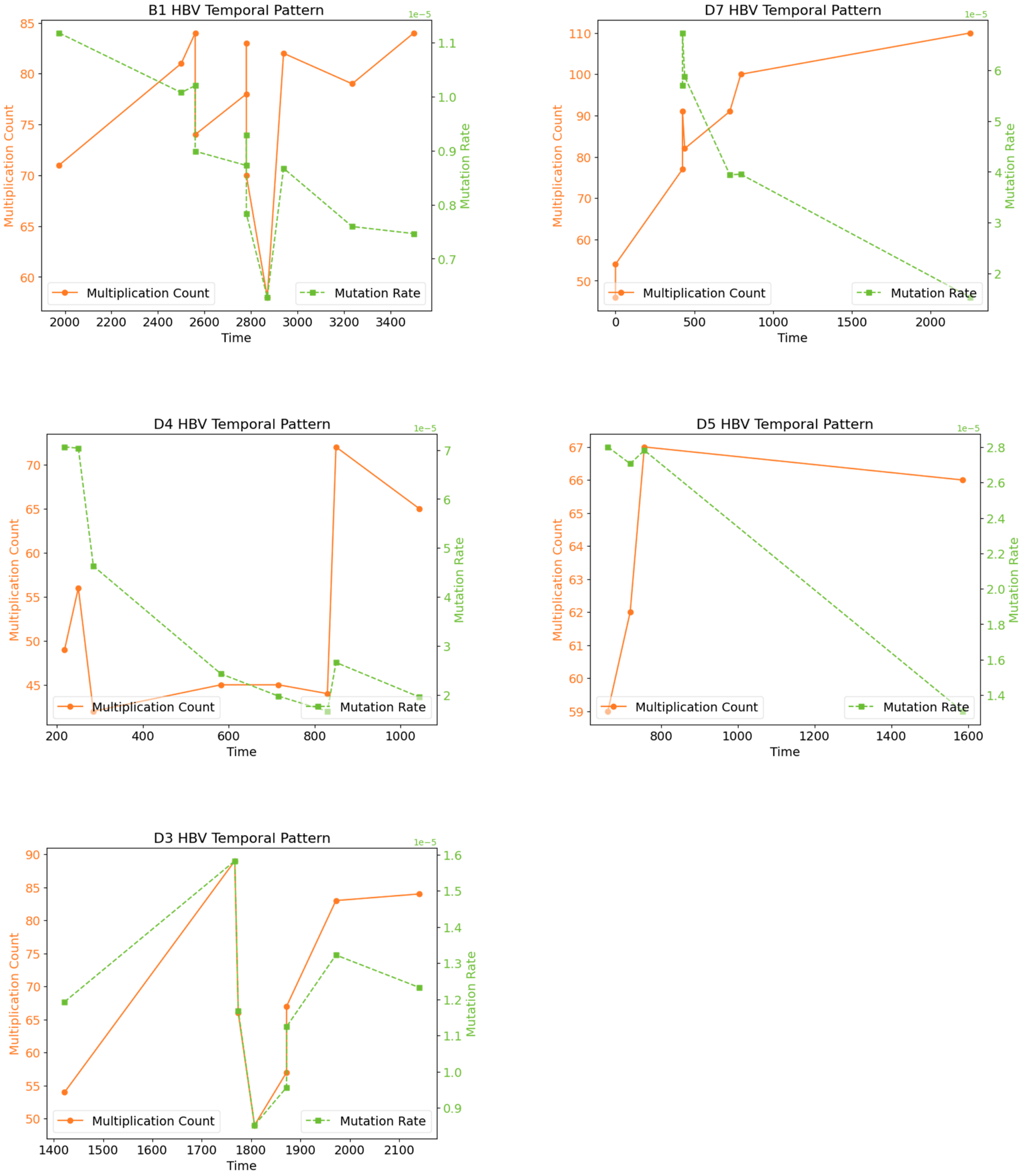


**Fig. S4.** Temporal mutation patterns of genotypes B and D HBV. Using the oldest data as a reference, the mutation count (yellow) and mutation rate (green) for HBV at different time points are statistically analyzed.

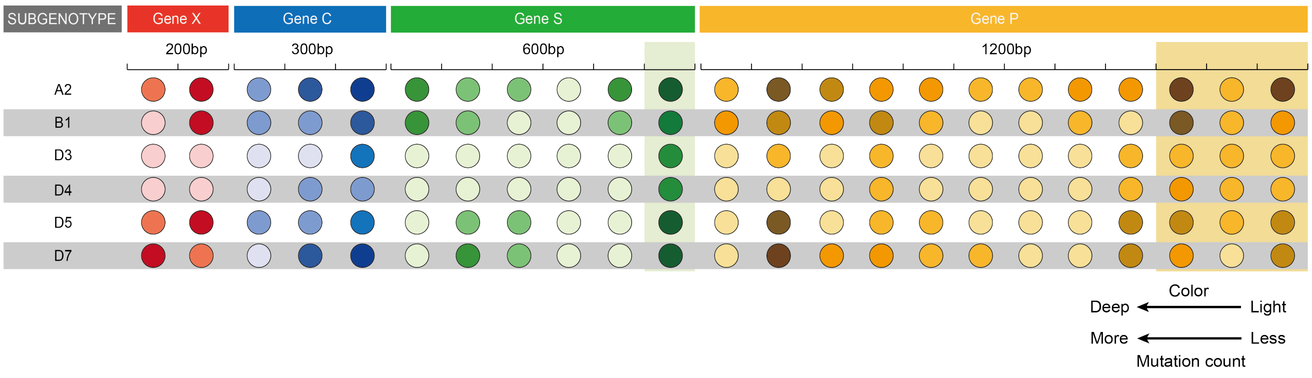


**Fig. S5.** Common mutation sites of subgenotypes.

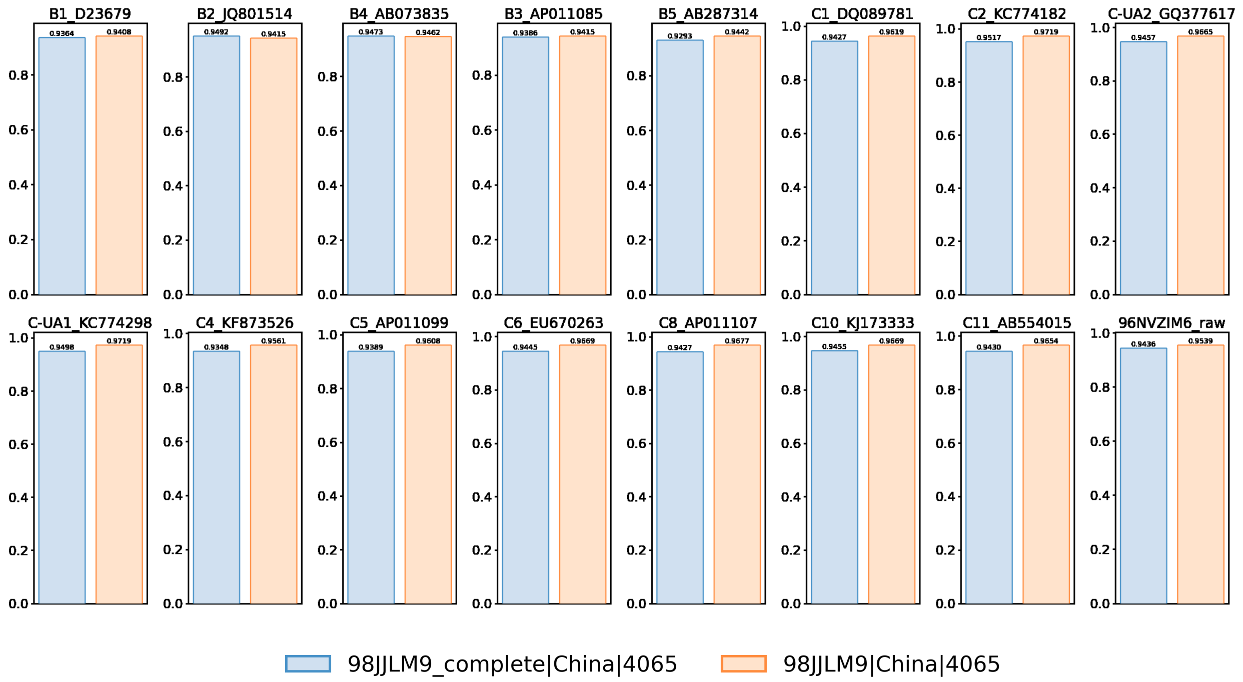


**Fig. S6.** Sequence identity for the imputed and raw sequences of 98JJLM9. In each subplot, the blue bar represents the imputed sequence, while the orange bar represents the raw sequence. Each subplot corresponds to a different subgenotype, as indicated by the title above each plot. The values on top of the bars indicate the result of sequence identity.

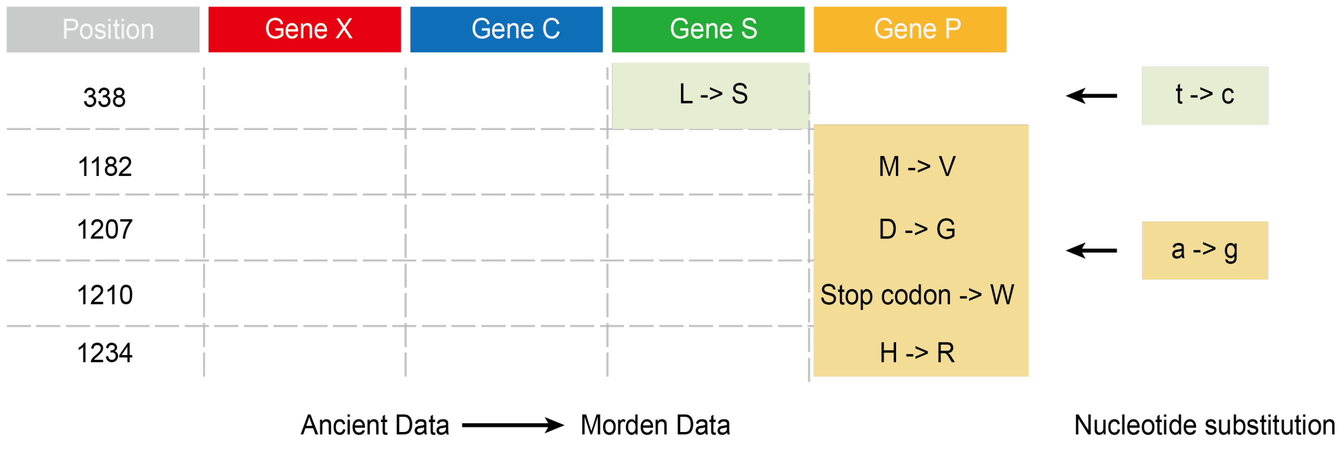


**Fig. S7.** Conserved mutations found in genotype A. A total of 1 mutation was found in the S gene and 4 mutations were found in the P gene, all of which were caused by the replacement of nucleotide "a" with nucleotide "g".
